# Telescope: Characterization of the retrotranscriptome by accurate estimation of transposable element expression

**DOI:** 10.1101/398172

**Authors:** Matthew L. Bendall, Miguel de Mulder, Luis Pedro Iñiguez, Aarón Lecanda-Sánchez, Marcos Pérez-Losada, Mario A. Ostrowski, R. Brad Jones, Lubbertus C. F. Mulder, Gustavo Reyes-Terán, Keith A. Crandall, Christopher E. Ormsby, Douglas F. Nixon

## Abstract

Characterization of Human Endogenous Retrovirus (HERV) expression within the transcriptomic landscape using RNA-seq is complicated by uncertainty in fragment assignment because of sequence similarity. We present Telescope, a computational software tool that provides accurate estimation of transposable element expression (retrotranscriptome) resolved to specific genomic locations. Telescope directly addresses uncertainty in fragment assignment by reassigning ambiguously mapped fragments to the most probable source transcript as determined within a Bayesian statistical model. We demonstrate the utility of our approach through single locus analysis of HERV expression in 13 ENCODE cell types. When examined at this resolution, we find that the magnitude and breadth of the retrotranscriptome can be vastly different among cell types. Furthermore, our approach is robust to differences in sequencing technology, and demonstrates that the retrotranscriptome has potential to be used for cell type identification. Telescope performs highly accurate quantification of the retrotranscriptomic landscape in RNA-seq experiments, revealing a differential complexity in the transposable element biology of complex systems not previously observed. Telescope is available at github.com/mlbendall/telescope.

**Author Summary:** Almost half of the human genome is composed of Transposable elements (TEs), but their contribution to the transcriptome, their cell-type specific expression patterns, and their role in disease remains poorly understood. Recent studies have found many elements to be actively expressed and involved in key cellular processes. For example, human endogenous retroviruses (HERVs) are reported to be involved in human embryonic stem cell differentiation. Discovering which exact HERVs are differentially expressed in RNA-seq data would be a major advance in understanding such processes. However, because HERVs have a high level of sequence similarity it is hard to identify which exact HERV is differentially expressed. To solve this problem, we developed a computer program which addressed uncertainty in fragment assignment by reassigning ambiguously mapped fragments to the most probable source transcript as determined within a Bayesian statistical model. We call this program, “Telescope”. We then used Telescope to identify HERV expression in 13 well-studied cell types from the ENCODE consortium and found that different cell types could be characterized by enrichment for different HERV families, and for locus specific expression. We also showed that Telescope performed better than other methods currently used to determine TE expression. The use of this computational tool to examine new and existing RNA-seq data sets may lead to new understanding of the roles of TEs in health and disease.

## Introduction

Transposable elements (TEs) represent the largest class of biochemically functional DNA elements in mammalian genomes(Dunham et al. 2012; Kellis et al. 2014) comprising nearly 50% of the human genome. As many of these transcriptionally active elements originated as retroelements, we refer to the set of RNA molecules transcribed from these elements in a population of cells as the retrotranscriptome. The contribution of the retrotranscriptome to the total transcriptome, cell-type specific expression patterns, and the role of retroelement transcripts in disease remain poorly understood (Magiorkinis et al. 2013). Although most TEs are hypothesized to be transcriptionally silent (due to accumulated mutations), recent studies have found many elements to be actively expressed and involved in key cellular processes. For example, aberrant expression of LINE-1 (L1) elements, the most expansive group of TEs, has been implicated in the pathogenesis of cancer (Wang-Johanning et al. 2003; Tang et al. 2017; Rodić et al. 2015; Ardeljan et al. 2017), while human endogenous retroviruses (HERVs) are reported to be involved in human embryonic stem cell differentiation(Grow et al. 2015; Göke et al. 2015) and in the pathogenesis of amyotrophic lateral sclerosis(Li et al. 2015). We, and others, have shown that HIV-1 infection increases HERV transcription(Garrison et al. 2007; Jones et al. 2012; Ormsby et al. 2012; Contreras-Galindo et al. 2012; Gonzalez-Hernandez et al. 2014). These lines of evidence therefore indicate that TEs have important roles in the regulation of human health and disease.

The ability to observe and quantify TE expression, especially the specific genomic locations of active elements, is crucial for understanding the molecular basis underlying a wide range of conditions and diseases(Flockerzi et al. 2008). Traditional techniques for interrogating the TE transcriptome include quantitative PCR (Muradrasoli et al. 2006; Rangwala et al. 2009) and RNA expression microarrays (Seifarth et al. 2003; Pérot et al. 2012; Gnanakkan et al. 2013; Young et al. 2014; Becker et al. 2017). However, these techniques are unable to discover elements not specifically targeted by the assay, and may fail to detect rare, previously unknown, or weakly expressed transcripts. High-throughput RNA sequencing (RNA-seq) promises to overcome many of these shortcomings, enabling highly sensitive detection of transcripts across a wide dynamic range. Mathematical and computational approaches for transcriptome quantification using RNA-seq are well established ((Mortazavi et al. 2008; Marioni et al. 2008), see review (Garber et al. 2011)) and provide researchers with reproducible analytical pipelines (Trapnell et al. 2010, 2012). Such approaches are highly effective at quantifying transcripts when sequenced fragments can be uniquely aligned to the reference genome, since the original genomic template for each transcript can be unambiguously identified (Trapnell et al. 2013; Conesa et al. 2016). In contrast, sequencing fragments generated by TEs often have high scoring alignments to many genomic locations with similar sequences, leading to uncertainty in transcript count estimates. Approaches that fail to account for these uncertainties may incorrectly estimate TE abundance and falsely detect significant changes in expression (Trapnell et al. 2013).

A growing number of studies are using high-throughput sequencing to characterize the retrotranscriptome. Three general approaches are used to deal with challenges of aligning short sequencing reads to repetitive elements. i) “Family-level” approaches combine read counts across all instances of a TE family, since fragments mapping to multiple genomic locations can often be uniquely assigned to a single repeat family. ii) “Heuristic” approaches simplify the problem of multi-mapped fragments by either discarding ambiguous reads (unique counts) or randomly assigning ambiguous reads to one of its best scoring alignments (best counts). Finally, iii) “statistical” approaches estimate the most probable assignment of fragments given a statistical model. Our approach, Telescope, implements a Bayesian statistical model for reassigning ambiguous fragments; previous work that has used statistical approaches include the TETranscripts package (Jin et al. 2015) and an ad hoc model implemented by (Santoni et al. 2012).

Here, we introduce Telescope, a tool which provides accurate estimation of TE expression resolved to specific genomic locations. Our approach directly addresses uncertainty in fragment assignment by reassigning ambiguously mapped fragments to the most probable source transcript as determined within a Bayesian statistical model. We implement our approach using a descriptive statistical model of the RNA-seq process and use an iterative algorithm to optimize model parameters. We use Telescope to investigate the expression of HERVs in cell types from the ENCODE consortium.

## Results

### Telescope: Single locus resolution of transposable element expression

Resolution of transposable element (including those of human endogenous retroviruses, HERVs) expression from RNA-seq data sets has been complicated by the many similarities of these repetitive elements. Telescope is a computational pipeline program that solves the problem of ambiguously aligned fragments by assigning each sequenced fragment to its most likely transcript of origin. We assume that the number of fragments generated by a transcript is proportional to the amount of transcript present in the sample; thus, the most likely source template for a randomly selected fragment is a function of its alignment uncertainty and the relative transcript abundances. Telescope describes this relationship using a Bayesian mixture model where the estimated parameters include the relative transcript abundances and the latent variables define the possible source templates for each fragment (Francis et al. 2013).

The first step in this approach is to independently align each fragment to the reference genome; the alignment method should search for multiple valid alignments for each fragment and report all alignments that meet or exceed a minimum score threshold (Fig 1A). Next, alignments are tested for overlap with known TE transcripts; transcript assignments for each fragment are weighted by the score of the corresponding alignment (Fig 1B and 1C). In our test cases, we typically find that less than 50% of the fragments aligning to TEs can be uniquely assigned to a single genomic location and many fragments have more than 20 possible originating transcripts.

**Fig 1.**
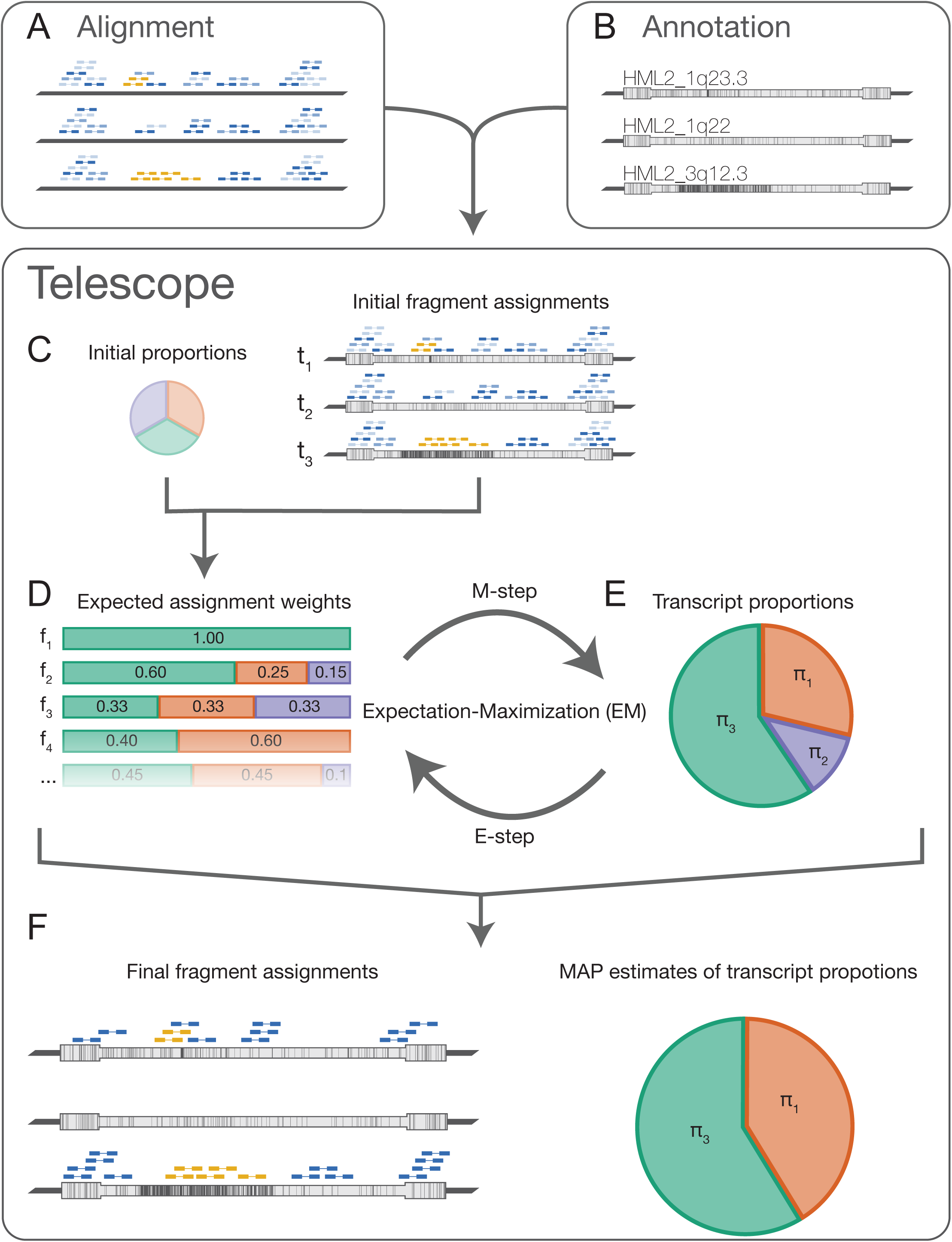
Telescope conceptual overview. A set of possible genomic locations for each fragment is determined by alignment to the reference genome. In order to find as many high-scoring mappings as possible, we use sensitive local alignment parameters and search for up to 100 alignments for each fragment using bowtie2 (A). Using an annotation containing known TE locations, Telescope intersects the aligned fragments with annotated TE loci (B,C). The set of alignments and corresponding alignment scores for each fragment are used to calculate the expected assignment weights, initially assuming equal expression for all elements (D). The assignment weights estimated in (D) are used to find the maximum likelihood estimate (MLE) for the proportion of each transcript (E). Next, we update the expected assignment weights, now assuming that the MLE represents our best estimate of transcript expression (D,E). The steps in panels (D) and (E) describe an expectation-maximization procedure, and we further refine the assignment weights and MLE by iterating until parameter estimates converge. Telescope produces a report that includes the maximum a posteriori estimate of the transcript proportions and the final number of fragments assigned to each transcript, as well as an updated alignment including the final fragment assignments (F).

Telescope estimates the transcript proportions and expected source templates using an expectation-maximization algorithm. In the expectation step (E-step), the expected value of the source template for each fragment is calculated under current estimates of transcript abundance (Fig 1D). The maximization step (M-step) finds maximum *a posteriori* estimates of the transcript abundance dependent on the expected values from the E-step (Fig 1E). These steps are repeated until parameter estimates converge (Fig 1D and 1E). Telescope reports the proportion of fragments generated by each transcript and the expected transcript of origin for each fragment (Fig 4F). The final counts estimated by Telescope correspond to actual observations of sequenced fragments and are suitable for normalization and differential analysis by a variety of methods. The software also provides an updated alignment with final fragment assignments that can be examined using common genome visualization tools. Telescope is available at github.com/mlbendall/telescope.

### Determination of HERV expression in major cell types from the ENCODE consortium

To investigate HERV expression in a robust way across a diverse platform of cell types we relied on publicly available RNA-seq data. The ENCODE data project is an invaluable source of genomic data from disparate sources and provides the opportunity to mine the transposable element expression in a setting of maximum genomic information. We profiled 13 human cell types, including common lines designated by the ENCODE consortium, as well as primary cell types, and applied our approach to determine HERV expression across the spectrum of human cell types, including normal or transformed, and contrasting cell lines with primary cells (Table 1).

**Table 1.** ENCODE cell types used in this study

| Cell Type | Description | Karyotype | Lineage | Tissue | Replicates |
| --- | --- | --- | --- | --- | --- |
| H1-hESC | Embryonic stem cell | Normal | ICM | ESC | 4 |
| GM12878 | B-lymphocyte | Normal | mesoderm | blood | 4 |
| K562 | Leukemia | Cancer | mesoderm | blood | 3 |
| HeLa-S3 | Cervical carcinoma | Cancer | ectoderm | cervix | 3 |
| HepG2 | Hepatocellular carcinoma | Cancer | endoderm | liver | 3 |
| HUVEC | Umbilical vein endothelial cells | Normal | mesoderm | vessel | 3 |
| SK-N-SH | Neuroblastoma | Cancer | ectoderm | brain | 1 |
| IMR90 | Fetal lung fibroblasts | Normal | endoderm | lung | 1 |
| A549 | Lung carcinoma | Cancer | endoderm | lung | 1 |
| MCF-7 | Mammary gland adenocarcinoma | Cancer | ectoderm | breast | 2 |
| CD20+ | CD20+ B cells | Normal | mesoderm | blood | 1 |
| CD14+ | CD14+ Monocytes | Normal | mesoderm | blood | 1 |
| NHEK | Epidermal keratinocytes | Normal | ectoderm | skin | 3 |

Over 2.7 billion sequenced fragments aligned to human reference hg38 with between 23.6% and 46.1% of the fragments in each sample aligning ambiguously to multiple genomic locations. Telescope intersected the aligned fragments with a set of 14,968 manually curated HERV loci belonging to 60 families (see methods) and identified over 27 million fragments that appear to originate from HERV proviruses. Most (80.1%) of these fragments aligned to multiple genomic locations; we used Telescope to reassign ambiguous fragments to the most likely transcript of origin and estimate expression at specific HERV loci.

We developed genome-wide maps of HERV expression for 8 of the analyzed cell types that had replicates (Table 1), and used CIRCOS (Krzywinski 2009) to visualize the data (Fig 2). The outer track is a bar chart showing the number of HERV loci in 10 Mbp windows, with the red part of the bar representing the number of loci that are expressed in one or more cell types. The 8 inner rings show the expression levels (log2 counts per million (CPM)) of 1365 HERV loci that were expressed at least one of the cell types examined. Moving from the outer ring to the inner ring are replicates for each of the 8 cell types with duplicates: H1-hESC, GM12878, K562, HeLa-S3, HepG2, HUVEC, MCF-7, and NHEK.

**Fig 2.**
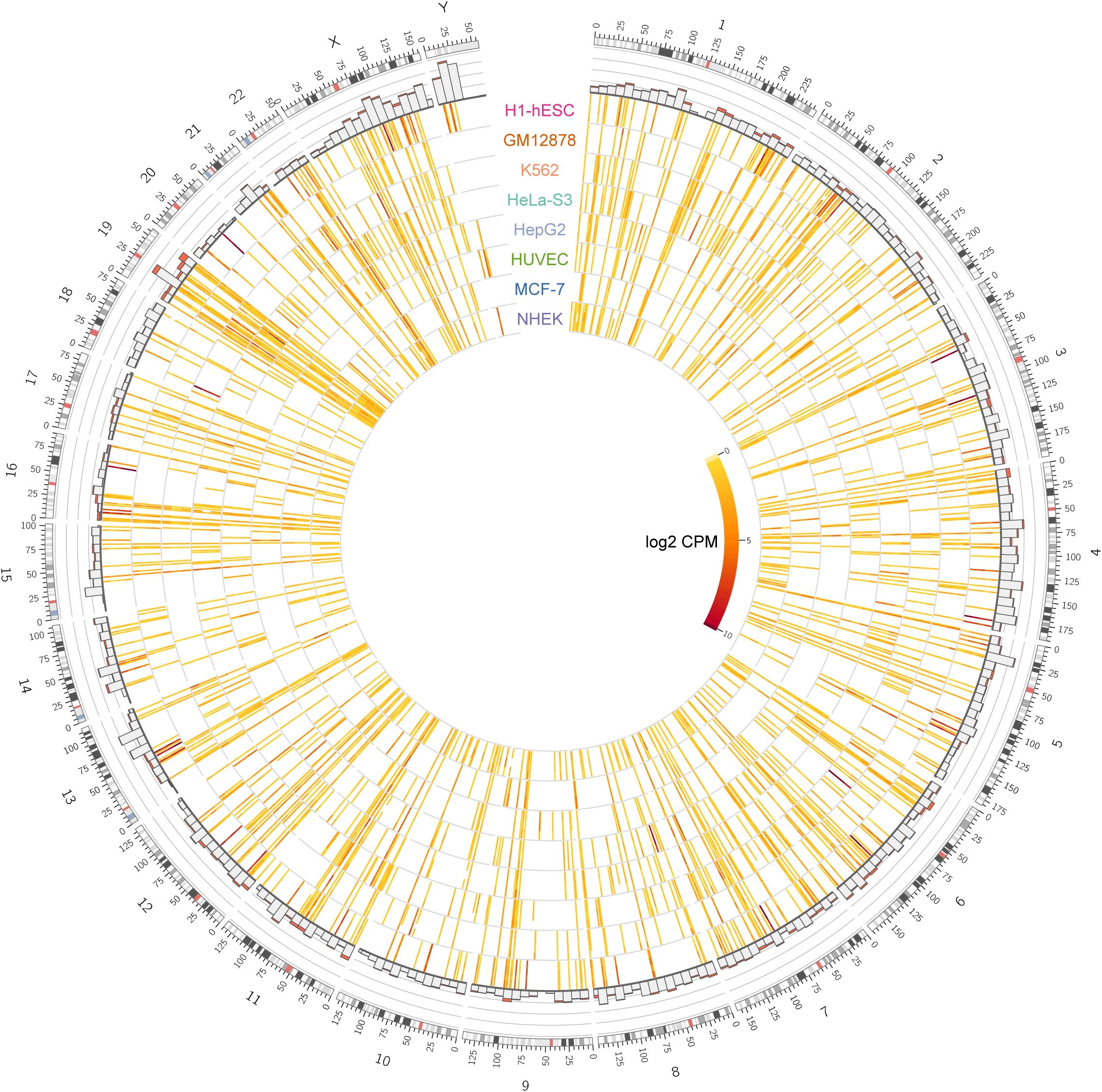
Genome-wide maps of locus-specific HERV expression for 8 ENCODE tier 1 and 2 cell types. The outer track is a bar chart showing the number of HERV loci in 10 Mbp windows, with the red part of the bar representing the number of loci that are expressed in one or more cell types. The 8 inner rings show the expression levels (log2 counts per million (CPM)) of 1365 HERV loci that were expressed at least one of the cell types examined. Moving from the outer ring to the inner ring are replicates for each of the 8 cell types with duplicates: H1-hESC, GM12878, K562, HeLa-S3, HepG2, HUVEC, MCF-7, and NHEK.

We found 1365 HERV loci that were expressed in at least one of the cell types (CPM > 0.5). Not all HERVs were expressed in all cell types, some were widely expressed in all cells, whereas others were only expressed in one or more cell type (Fig 2). There is also a spectrum of differential HERV expression, with some HERVs having significantly higher expression than others. On a chromosome by chromosome analysis, there are certain regions of the genome that have minimal HERV expression, while other regions appear dense in HERV expression. There are areas of scarce HERV expression on chromosomes 3, 5, 9, 15, and the Y chromosome (Fig 2). Interestingly, the Y chromosome is host to a greater density of HERV locations, yet they are mostly silent. In contrast, several chromosomes exhibit a greater than expected number of active HERV locations, i.e. chromosome 19 (S1 Fig) and chromosome 6 (S2 Fig).

### HERV Locus-specific-analysis

To ascertain, global, family and locus level specific HERV expression, we assessed the number of HERVs expressed in each cell type. All cell types expressed HERVs; the number of expressed loci ranged from 216 (in MCF-7), to 533 (H1-hESC) (Fig 3A). The number and proportion of cell type specific locations (expressed in only one cell) differed among cell types. Nearly half (46.3%) of locations expressed in H1-hESC were not expressed in any other cell type, while 89.3% of locations expressed in MCF-7 were also present in other cell types (Fig 3A). This suggests that regulatory networks are shared among some cell types but not others. We next examined the relative contribution of HERV families to overall HERV transcription and found that different cell types could be characterized by enrichment for different HERV families. For example, HERVH accounted for 91.8% of the transcriptomic output in H1-hESC cells, while HERVE was dominant in K562 cells (24.4%) (Fig 4A). Other families, such as HERVL, were evenly distributed across cell types, both in number of expressed locations and in expression levels (Fig 4B). Resolving the most highly expressed specific locations in each cell type at a locus specific level shows that the distribution of expression varies among cell types. (Fig 3C). For example, HepG2 is characterized by high expression from a single locus, while H1-hESC has many locations that are activated.

**Fig 3.**
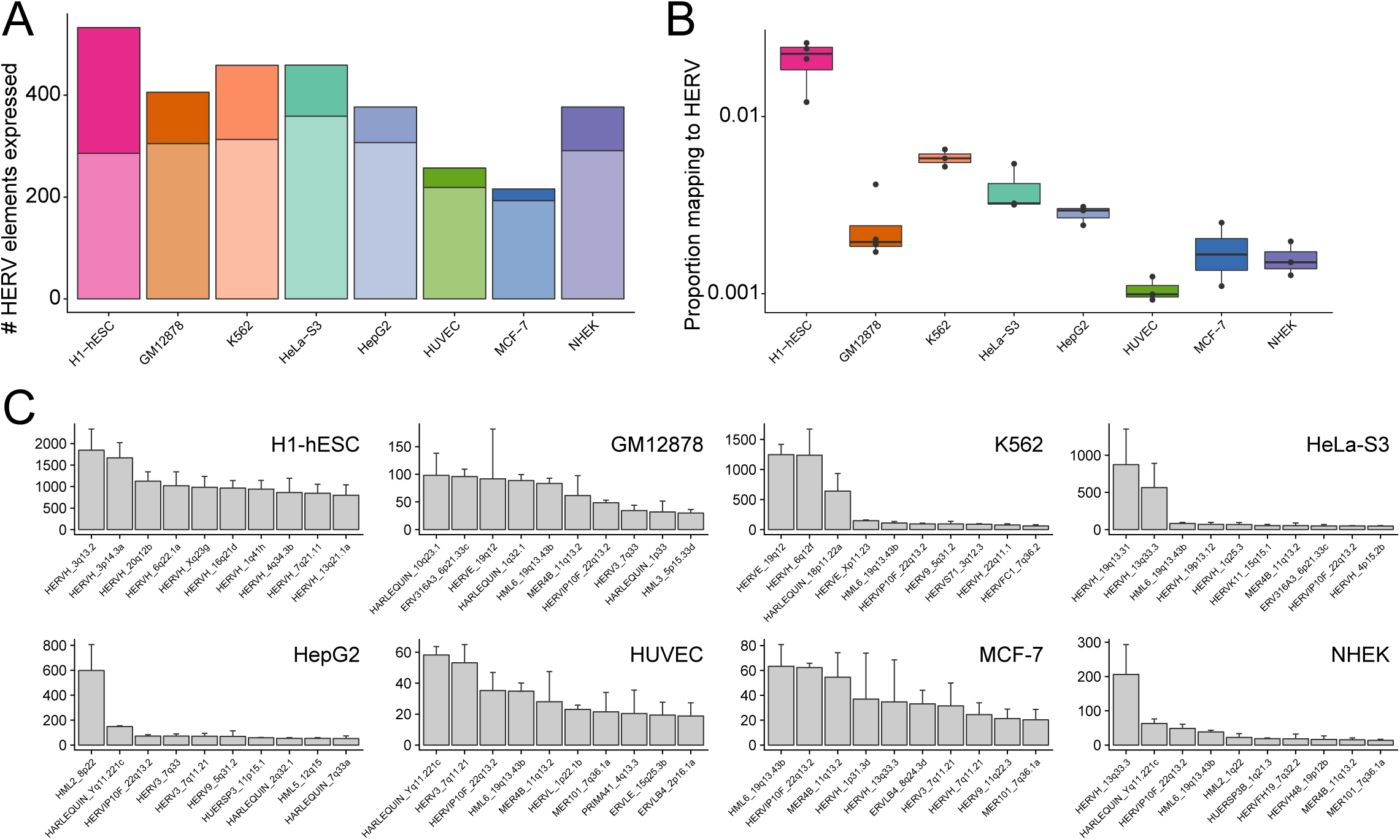
Overall HERV expression patterns. (A) Number of HERV elements that are expressed for each cell type; expressed loci have CPM > 0.5 in the majority of replicates. The darker section of the bar corresponds to expressed loci that are unique to cell type, while the lighter part is expressed in other cell types. (B) The proportion of mapped RNA-seq fragments that are generated from HERV transcripts in each of eight replicated cell types. Each point is one replicate; boxplot shows the median and first and third quartiles. (C) Top 10 most highly expressed loci for each cell type. Height of the bar is average CPM of all replicates with error bars representing the standard error calculated from replicates CPM values.

**Fig 4.**
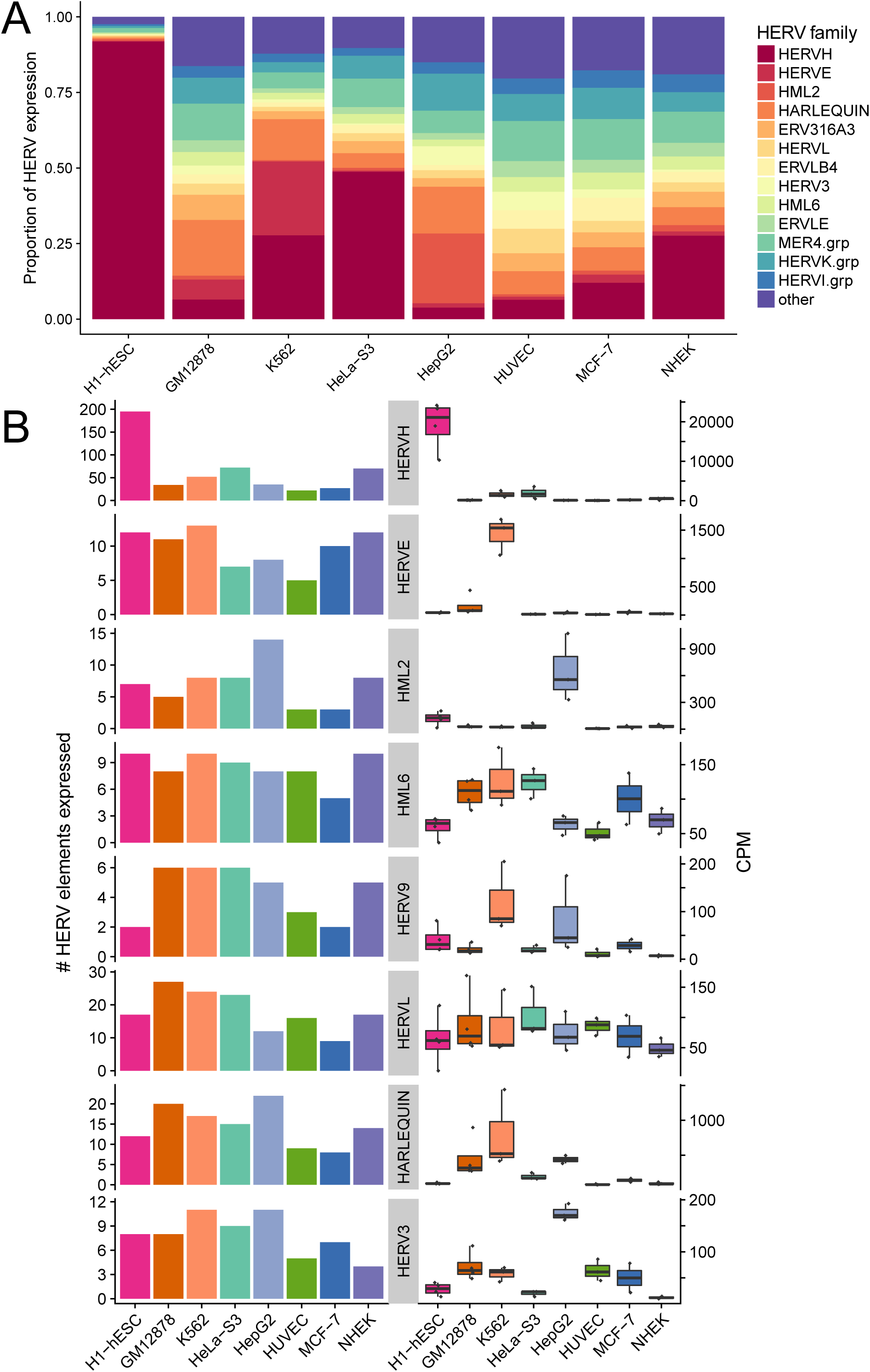
Family-level HERV expression profiles using Telescope. Family-level HERV expression profiles were computed from locus-specific profiles (generated by Telescope) by summing expression across all locations within each family. (A) The proportion of fragments assigned to each HERV family relative to the total amount of HERV expression. Families that account for at least 5% of total HERV expression in at least one cell type are shown, with the remaining families in “other”. (B) Number of expressed HERV loci and fragment counts per million mapped fragments (CPM) for selected HERV families.

### HERV expression profiles generated by Telescope are cell type specific

Previous work has suggested that estimates of HERV expression are highly sensitive to sequencing technology used, and differences due to sequencing technology can obscure biological differences due to cell type (Haase et al. 2015). Since aligning shorter fragments (i.e. single-end reads) tends to produce more ambiguously mapping fragments compared to longer fragments, we hypothesized that Telescope (which resolves ambiguity) would create HERV expression profiles that are robust to differences in sequencing technology. Hierarchical clustering of all 30 polyA RNA-seq HERV profiles shows that replicates from the same cell type cluster most closely with other samples from the same cell type, regardless of the sequencing technology used (Fig 5A). Clusters for all cell types had significant support using multiscale bootstrap resampling (approximately unbiased (AU) > 95%). Principal component analysis (PCA) also indicates that cell type, not sequencing technology, is associated with the strongest differences among expression profiles. The first principal component, accounting for 44% of the total variance in the data, separates H1-hESC samples from all other samples (Fig 5B). The second and third components further separate the samples into the other 12 cell types, and capture 13% and 10% of the total variance, respectively. Interestingly, the second component separates blood-derived cell types (K562, GM12878, CD20+ and CD14+) from the other cell types, suggesting that cells derived from the same tissue may share similarities in HERV expression profiles.

**Fig 5.**
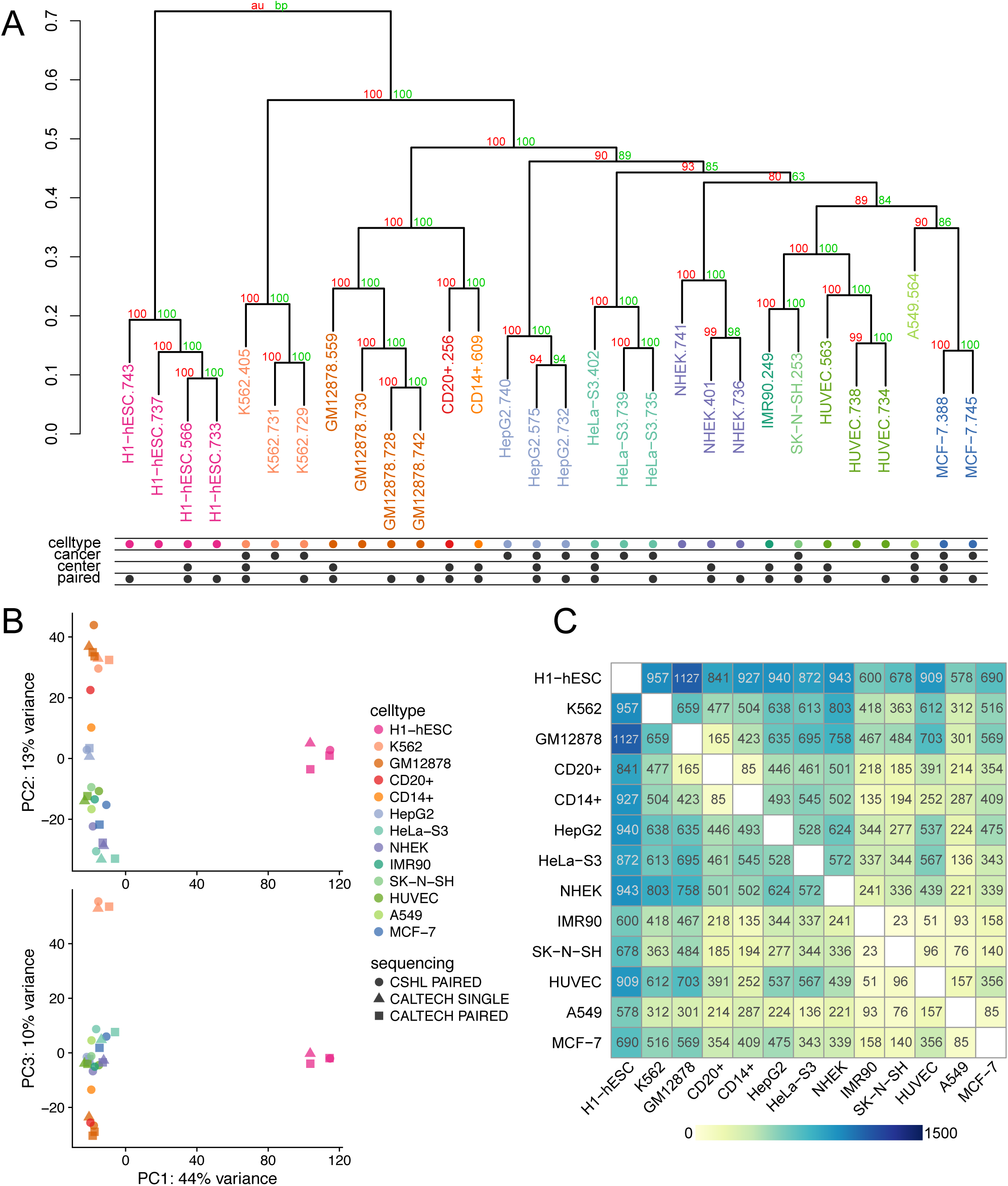
Cell type characterization based on HERV expression profiles using unsupervised learning and linear models. Unsupervised learning and linear modeling were used to identify patterns in HERV expression profiles generated by Telescope for 30 polyA RNA-seq datasets from 13 cell types. (A) Similarities among normalized expression profiles were explored using hierarchical cluster analysis. Supporting p-values were based on 1000 multiscale bootstrap replicates and calculated using Approximately Unbiased (AU, red) and Bootstrap probability (BP, green) approaches. (B) Principal component analysis (PCA) of normalized expression profiles. The first component accounts for 44% of the variance in the data, and is plotted against component 2 and 3, which account for 13% and 10% of the variance, respectively. (C) Heatmap of the number of HERV elements found to be significantly differentially expressed (DE) among each pair of cell types. Significance was determined using cutoffs for the false discovery rate (FDR < 0.1) and log2 fold change (abs(LFC) > 1.0). Yellow indicates low numbers of differentially expressed elements, while blue indicates high numbers.

We further explored differences among cell types using differential expression (DE) analysis. Pairwise contrasts between cell types were performed to determine the number of significant DE loci (FDR < 0.1, abs(LFC) > 1.0) (Fig 5C). As found in the unsupervised analysis, HERV expression in H1-hESC was drastically different from other cell types, with between 578 and 1127 significantly DE loci.

Finally, we asked whether heuristic approaches for TE quantification would be sufficient to identify cell type specific signal in the data or whether these approaches would be sensitive to other variables. We performed the same unsupervised analyses with HERV expression profiles obtained using unique and best counting approaches. Hierarchical clustering using unique count expression profiles produced a very similar topology to that found using Telescope (S3 Fig). Replicate samples were properly clustered by cell type, though GM12878 and HUVEC had slightly less bootstrap support. In contrast, clustering with the best count profiles did not recover all cell type clusters; two HeLa-S3 samples clustered with H1-hESC, while the third was more similar to A549 cells (S3 Fig). There was also less support for several clusters, including one cell type cluster (NHEK) that did not meet the 95% threshold.

### Performance of Telescope compared to current methods

In order to examine the sensitivity and biases of computational approaches for quantifying TE expression, we designed simulation experiments with known expression values. Earlier studies have suggested that the HERV-K(HML-2) family (hereafter referred to as HML-2) is expressed in human tissue and may be relevant to human health (Hohn et al. 2013; Grow et al. 2015; Li et al. 2015; Weiss 2016). Furthermore, its relatively few family members (∼90 distinct genomic loci (Subramanian et al. 2011)) and high nucleotide identity make HML-2 a good model for studying TE expression. Here, we report on the performance of each method to detect locus-specific expression of HML-2 by simulating RNA-seq fragments. We simulated 25 independent datasets, each simulation consisted of 10 randomly chosen HML-2 loci and a fragment count, which could be interpreted as an expression value. We used the following TE quantification approaches for estimating locus specific HML-2 expression: unique counts, best counts, RepEnrich (Criscione et al. 2014), TETranscripts (Jin et al. 2015) and Telescope.

The greatest strength of the unique counts approach was the low false detection rate since across all 25 simulations (41.5K simulated fragments), only 6 fragments were incorrectly assigned. However, unique counts consistently underestimated expression levels with ∼60% of all estimates (151 out of 250) missing at least 50% of the true expression (Fig 6A). One striking example of this underestimation was for HML2_5q33.3; this locus did not generate any fragment that could be counted by unique counts despite being expressed in 5 simulations. We presume that the underestimation is a direct consequence of discarding ambiguously mapped reads, as the unique counts discarded 62.6% of the simulated fragments.

**Fig 6.**
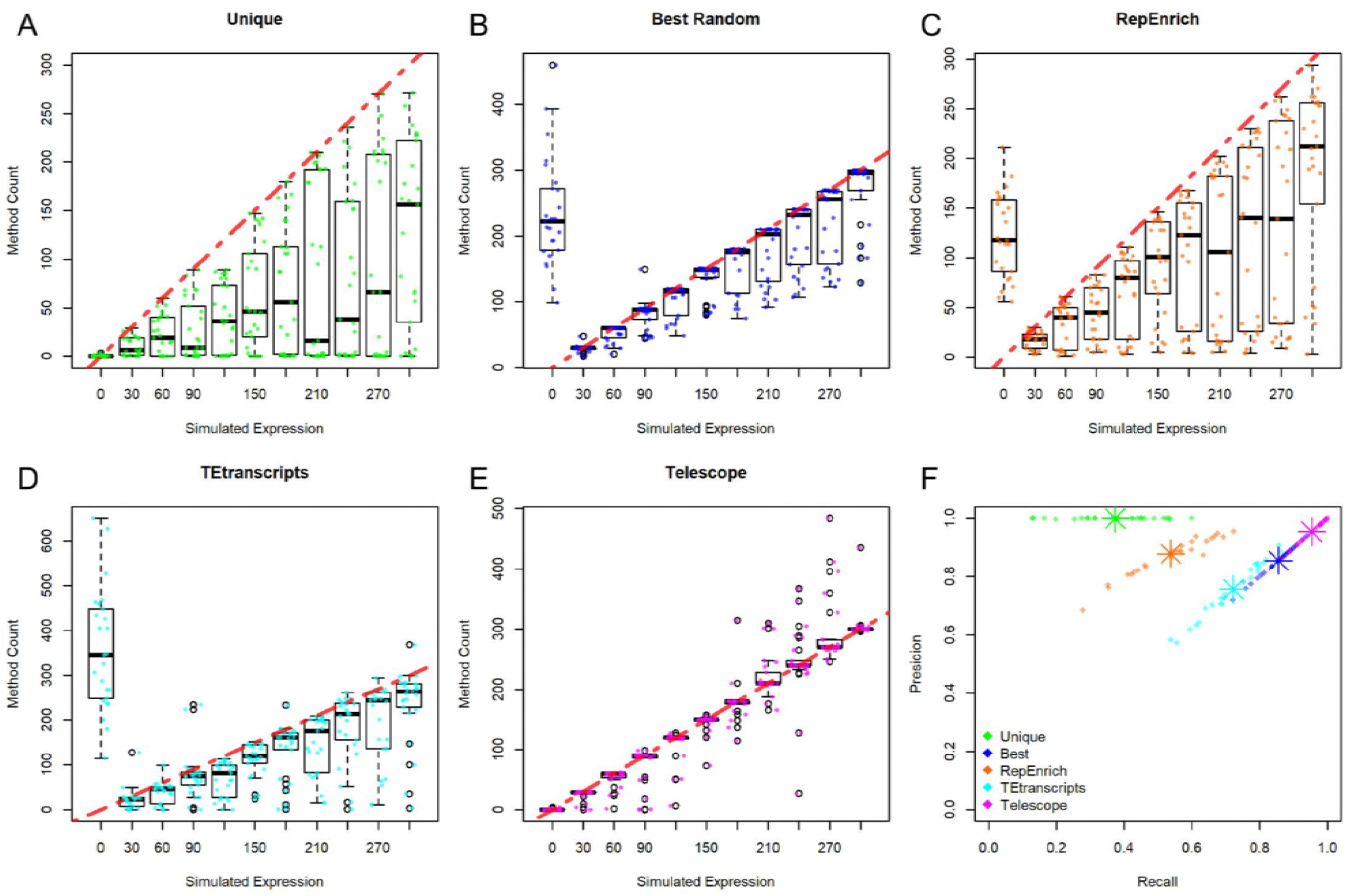
Comparison of performance results for unique counts, best counts, RepEnrich, TEtranscripts and Telescope using simulated data. 25 RNA-seq samples were simulated, each sample consisted of 10 randomly chosen HML-2 loci with each having different expression value. The possible expression values are shown along the x-axis, 0 represent all HML-2 not expressed, red dashed line represents the expected expression value. A box plot representing the count distribution from each expression value is plotted. The resulted count from the different counting method from each simulated expression value per sample is plotted over the boxplot. Counting methods tested: (A) unique count, (B) best count, (C) RepEnrich, (D) TEtranscript, (E) Telescope. (F) The precision and recall for each sample simulated as well as the mean of both are shown for all methods.

In contrast to unique counts, the best counts approach offers greater sensitivity (Fig 5F). Instead of discarding ambiguously mapped fragments, all the fragments are used, resulting in more accurate expression estimates (Fig 5B). The majority of fragments were assigned to the true source transcript, while incorrect hits account for 14.2% of the total fragments. These off-target assignments resulted in false detection of unexpressed loci in each simulation, representing a major drawback of this approach (Fig 5B). Despite the high sensitivity of best counts, we conclude that the high number of incorrect detections outweighs the possible advantages of using this approach.

In order to make a direct comparison between Telescope and RepEnrich, which quantifies TE families at the family level, we modified RepEnrich annotations to give each locus a unique “family” name. We found that expression was underestimated by this approach for all expressed HML-2 elements (Fig 5C). We attribute this bias to the large number of fragments that were discarded by this approach, 38.8%. Nevertheless, the number of discarded reads is fewer than the unique counts approach where 62.6% of all fragments were omitted. Another negative aspect of RepEnrich is that 12.1 % of the counted fragments were assigned to non-expressed TE.

We tested TETranscripts on our simulated data with the TE counting method set under “multi” mode. We provided the algorithm with our HERV annotation thus results could be compared with Telescope. TETranscripts showed a better performance than RepEnrich as shown in Jin *et al.* (2015) involving multiple mapped fragments assignment but 21.7% of all fragments were assigned incorrectly to a non-expressed HML-2. Based on the precision and recall of all methods tested TETranscripts performed as the third best TE single locus counting method on the simulated data (S4 Fig).

Finally, we tested Telescope’s ability to reassign ambiguous alignments and estimate locus-specific fragment counts using the same simulation data. On average 57.8% of the simulated fragments aligned to multiple loci, which need to be reassigned. Telescope reassigned ambiguously mapped fragments to the expected transcript of origin according to our model (see methods) and reported the final number of fragments aligned to each TE. The estimated levels of expression, calculated with Telescope, from each HML-2 resembled closely the simulated levels (Fig 5E). Only 3 HML-2 loci that were simulated to be expressed did not present any fragments and 99.9% of all simulated fragments were counted with the Telescope approach.

Of all methods considered here, Telescope had the highest rate of precision and recall from all other counting methods tested (Fig 5F). In contrast to the best counts approach, the second best (S4 Fig), Telescope assigned only 15 fragments were assigned to genomic locations that were not expressed, while 5871 fragments were assigned incorrectly by best counts. Deviations of Telescope estimates from true expression levels, as measured by F1-score, was the highest of all approaches (S4 Fig). These simulation results demonstrate that Telescope resolves ambiguously aligned fragments and produces unbiased estimates of TE expression that are robust to sequencing error.

## DISCUSSION

High-throughput RNA sequencing has enabled the simultaneous characterization and quantification of an entire transcriptome with remarkable resolution and sensitivity. Current studies have primarily focused on protein-coding transcripts, with greater attention being given to non-coding and micro RNAs in recent years. The transposable elements represent another major biochemically active group of transcripts and are increasingly recognized as important regulators in complex biological systems and disease yet have been largely ignored in the literature. We present a novel software program, Telescope, that can be used to mine new or existing RNA-seq datasets to accurately quantify the expression of TEs. The key advantage of our approach is the capability to localize TE expression to an exact chromosomal location.

As TEs are repetitive elements located throughout the genome, existing programs have limitations in performing accurate alignments from RNA fragments because of sequence similarity. The management of alignment uncertainty has been approached in several ways. The unique count approach discards fragments that align ambiguously, but this approach underestimates or fails to detect gene expression. The best count method assigns each fragment to the source template with the best scoring alignment, but this underestimates TEs that are truly expressed and spuriously detects those that are in fact absent. While the family method of alignment mitigates uncertainty in alignments by classification according to repeat family, this method does not locate the genomic site of TE transcription. Our approach, Telescope, reassigns fragments to the most likely originating transcript using Bayesian mixture model that relating the relative transcript abundances to the possible source templates for each fragment. This approach thus resolves ambiguously aligned fragments and results in accurate quantification of TE loci for differential analysis.

Telescope will have widespread utility in other settings. Studies on TE expression have become prominent in studies of embryonic stem cell development (Grow et al. 2015)(Göke et al. 2015), neural cell plasticity (Muotri et al. 2010; Gage and Muotri 2012), oncogenesis (Wang-Johanning et al. 2003; Rakoff-Nahoum et al. 2006; Takahashi et al. 2008; Tang et al. 2017; Rodić et al. 2015; Ardeljan et al. 2017), psychiatric and neurological disorders(Perron et al. 2012; Christensen 2016; Mortelmans et al. 2016) and autoimmune diseases (Nexø et al. 2015; Hanke et al. 2016). As the breadth of knowledge on TEs expands, expression profiling of TEs using Telescope will allow scientists to discover unique and collective TE transcripts involved in the biology of complex systems.

## Methods

### Fragment reassignment mixture model

Telescope implements a generative model of RNA-seq relating the probability of observing a sequenced fragment to the proportions of fragments originating from each transcript. Formally, let *F* = [*f*_1_, *f*_2_,…, *f*_*N*_] be the set of *N* observed sequencing fragments. We assume these fragments originate from *K* annotated transcripts in the transcriptome *T* = [*t*_1_, *t*_2_,…, *t*_*K*_]. In practice, annotations fail to identify all possible transcripts that generate fragments, thus we include an additional category, *t*_0_, for fragments that cannot be assigned to annotated transcripts. Let *G* = [*G*_1_, *G*_2_,…, *G*_*N*_] represent the true generating transcripts for *F*, where *G*_*i*_ *∈ T* and *G*_*i*_ = *t*_*j*_ if *f*_*i*_ originates from *t*_*j*_. Since the process of generating *F* from *T* cannot be directly observed, the true generating transcripts *G* are considered to be “missing” data. The objective of our model is to estimate the proportions of *T* by learning the generating transcripts of *F*.

As described above, the alignment stage identifies one or more possible alignments for each fragment, along with corresponding alignment scores. Let *q*_*i*_ = [*q*_*i*0_, *q*_*i*1_,…, *q*_*ik*_] be the set of mapping qualities for fragment *f*_*i*_, where *q*_*ij*_ = Pr (*f*_*i*_ |*G*_*i*_ = *t*_*j*_) represents the conditional probability of observing *f*_*i*_ assuming it was generated from *t*_*j*_; we calculate this by scaling the raw alignment score by the maximum alignment score observed for the data. We write the likelihood of observing uniquely aligned fragment *f*_u_ as a function of the conditional probabilities *q*_u_ and the relative expression of each transcript for all possible generating transcripts *G*_u_

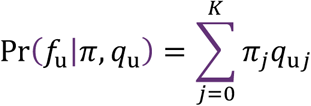

where ***π*** = [*π*_0_, *π*_1_,…, *π*_*K*_] represents the fraction of observed fragments originating from each transcript. Note that *q*_u*j*_ = 0 for all transcripts that are not aligned by *f*_u_. For non-unique fragments, we introduce an additional parameter in the above likelihood to reweight each ambiguous alignment among the set of possible alignments. The probability of observing ambiguous fragment *f*_a_ is given by

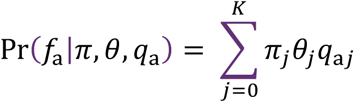

where ***θ*** = [*θ*_0_, *θ*_1_,…, *θ*_*k*_] is a reassignment parameter representing the fraction of non-unique reads generated by each transcript.

Using these probabilities of observing ambiguous and unique fragments, we formulate a mixture model describing the likelihood of the data given parameters ***π*** and ***θ***. The *K* mixture weights in the model are given by ***π***, the proportion of all fragments originating from each transcript. To account for uncertainty in the initial fragment assignments, let *x*_*i*_ = [*x*_*i*0_, *x*_*i*1_,…, *x*_*ik*_] be a set of partial assignment (or membership) weights for fragment *f*_*i*_, where 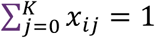 and *x*_*ij*_ = 0 if *f*_*i*_ does not align to *t*_*j*_. We assume that *x*_*i*_ is distributed according to a multinomial distribution with success probability ***π*.** Intuitively, *x*_*ij*_ represents our confidence that *f*_*i*_ was generated by transcript *t*_*j*_. In order to simplify our notation, we introduce an indicator variable ***y*** = [*Y*_1_, *Y*_2_,…, *Y*_*N*_] where *Y*_*i*_ = 1 if *f*_*i*_ is ambiguously aligned and *y*_*i*_ = 0 otherwise. The complete data likelihood is

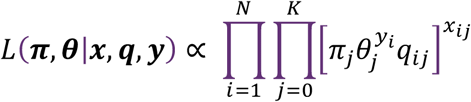

### Parameter estimation and fragment reassignment by EM

Telescope iteratively optimizes the likelihood function using an expectation-maximization algorithm (Dempster et al. 1977). First, the parameters ***π*** and ***θ*** are initialized by assigning equal weight to all transcripts. In the expectation step, we compute the expected values of *x*_*i*_ under current estimates of the model parameters. The expectation is given by the posterior probability of *x*_*i*_:

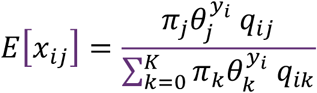

In the M-step we calculate the maximum a posteriori (MAP) estimates for ***π*** and ***θ***

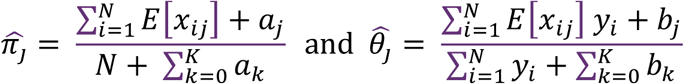

where *a*_*j*_ and *b*_*j*_ are prior information for transcript *t*_*j*_. Intuitively, these priors are equivalent to adding unique or ambiguous fragments to *t*_*j*_; providing non-zero values for these parameters prevents parameter estimates from converging to boundary values. Convergence of EM algorithms to local maxima has been shown by (Wu 1983), and is achieved when the absolute change in parameter estimates is less than a user defined level, typically *ε* < 0.001.

### HERV Annotations

A Telescope analysis requires an annotation that defines the transcriptional unit of each TE to be quantified. For HERV proviruses, the prototypical transcriptional unit contains an internal protein-coding region flanked by LTR regulatory regions. Existing annotations, such as those identified by RepeatMasker (Tarailo-Graovac and Chen 2009) (using the RepBase database (Jurka et al. 2005)) or Dfam (Wheeler et al. 2013) identify sequence regions belonging to TE families but do not seek to annotate transcriptional units. Both databases represent the internal region and corresponding LTRs using separate models, and the regions identified are sometimes discontinuous. Thus, a HERV transcriptional unit is likely to appear as a collection of nearby annotations from the same HERV family.

Transcriptional units for HERV proviruses were defined by combining RepeatMasker annotations belonging to the same HERV family that are located in adjacent or nearby genomic regions. Briefly, repeat families belonging to the same HERV family (internal region plus flanking LTRs) were identified using the RepBase database (Jurka et al. 2005). RepeatMasker annotations for each repeat family were downloaded using the UCSC table browser (Karolchik et al. 2004) and converted to GTF format, merging nearby annotations from the same repeat family. Next, LTR found flanking internal regions were identified and grouped using BEDtools (Quinlan and Hall 2010). HERV transcriptional units containing internal regions were assembled using custom python scripts. Each putative locus was categorized according to provirus organization; loci that did not conform to expected HERV organization or conflicted with other loci were visually inspected using IGV (Thorvaldsdóttir et al. 2013) and manually curated. As validation, we compared our annotations to the HERV-K(HML-2) annotations published by (Subramanian et al. 2011); the two annotations were concordant. Final annotations were output as GTF files and are available; all annotations, scripts, and supporting documentation are available at https://github.com/mlbendall/telescope_annotation_db.

### Simulated HML-2 expression data

We simulated 25 independent datasets, each consisted of randomly chosen 10 HML-2 which were expressed at different level, ranging from 30 to 300 fragments per locus. Using the expression pattern and the chosen HML-2, we simulated sequencing fragments with the Bioconductor package for RNA-seq simulation, Polyester (Frazee et al. 2014). All simulations used the parameters of read length: 75 bp; average fragment size: 250; fragment size standard deviation: 25; and an Illumina error model with an error rate of 5e-3.

### Alignment to reference genome

Sequenced fragments from each sample or simulation were aligned to human reference genome hg38 using bowtie2. Alignment options were specified to perform a sensitive local alignment search (--very-sensitive-local) with up to 100 alignments reported for each fragment pair (-k 100). The minimum alignment score threshold was chosen so that fragments with ∼95% or greater sequence identity would be reported (--score-min L,0,1.6).

### Software Availability

All scripts used for simulating and analyzing data are available at https://github.com/mlbendall/TelescopeEncode. The Telescope package is available at https://github.com/mlbendall/telescope.

## Author contributions

M.L.B., L.P.I, K.A.C. A.L.S., M.P.-L. and C.E.O developed the mathematics and statistics. L.P.I., D.G.H. and M.S. performed the experiments. M.M., and L.C.M designed the experiments and performed the analysis. M.L.B. implemented the software. M.L.B., M.M.R., M.A.O, R.B.J., G.R-T, K.A.C. and D.F.N. conceived the research. All authors wrote and approved the manuscript.

## Acknowledgments

The work was supported in part by US National Institutes of Health grants: CA206488 (DFN), AI076059 (DFN), UL1TR001876 (KAC), GM113886 (LCFM), and GM113886-01S1 (LCFM). MP-L was partially supported by DC CFAR pilot and CFAR 1P30AI117970 awards. MLB is a predoctoral student in the Systems Biology Program of the Institute for Biomedical Sciences at the George Washington University. This work is from a dissertation to be presented to the above program in partial fulfillment of the requirements for the Ph.D. degree.

We thank Timothy Powell, Rodrigo Duarte and Deepak Srivastava for constructive reading of the manuscript.

